# Experience-dependent tuning of early olfactory processing in the adult brain

**DOI:** 10.1101/558734

**Authors:** Christopher M. Jernigan, Rachael Halby, Richard C. Gerkin, Irina Sinakevitch, Fernando Locatelli, Brian H. Smith

## Abstract

Experience-dependent plasticity in the central nervous system allows an animal to adaptively change their responses to stimuli over different time scales. In this study we explored the different time frames and mechanisms over which olfactory experience-dependent plasticity works. We measured the impact of experience on early olfactory processing by comparing naturally foraging animals with a diverse olfactory experience to animals from the same cohort that experienced a chronic reduction in adult olfactory experience. We placed age-matched sets of full-sister honey bees, *Apis mellifera*, into two different olfactory conditions, in which animals were allowed to forage *ad libitum*. In one condition (T), we reduced the olfactory experience of foraging bees by placing them in a tent in which both sucrose and pollen resources were associated with a single odor. In the second condition (F), bees were allowed to forage freely and therefore receive a diversity of naturally occurring resource-associated olfactory experiences. We found that bees with a reduced olfactory experience had less developed antennal lobes when compared to experienced foragers, suggesting early-adult sensory experience influences the development of olfactory processing. We next measured the antennal lobe glomerular responses to odors using calcium imaging, and found that diverse olfactory experience of bees also enhances the inter-individual variation in the glomerular response profiles to odors. Last, we measured the impact of this treatment in an olfactory learning assay. We found that bees with a reduced olfactory experience had more difficulty picking an odor out of a mixture, which led them to generalize more (or respond similarly) to different mixture components than bees with richer olfactory experiences. Our study highlights the impact of individual experience at multiple levels (i.e., behavioral, physiological, developmental) on early olfactory processing.

## INTRODUCTION

Odors are encoded in primary olfactory processing neuropils—the olfactory bulb (OB) or antennal lobes (AL)—in the brains of animals as diverse as insects and mammals, including humans (Hildebrand and Shepherd, 1997, Sinakevitch et al., 2017). Canonically, in these brain regions, the combinatorial nature of activity patterns that encode different odors is conserved across animals and are species specific (Galizia et al., 1999b). These olfactory codes result from binding of volatile chemicals to peripheral olfactory receptor neurons (ORNs) that then project, via axon terminals, to glomeruli in the OB in mammals or the AL in insects (Buck and Axel, 1991, Hildebrand and Shepherd, 1997, Sinakevitch et al., 2017). Glomeruli are spherical, highly synapse-rich areas in the OB and AL. Odor processing in glomeruli is influenced by several different types of excitatory and inhibitory local interneurons (LNs) as well as by modulatory neurons that encode, among other things, reward (Farooqui et al., 2003, Hammer and Menzel, 1998, Ramaekers et al., 2001, Sachse and Galizia, 2002, Schäfer and Bicker, 1986, Shang et al., 2007, Sinakevitch et al., 2011, Sinakevitch et al., 2013, Sinakevitch et al., 2017). Projection neurons (PNs) or mitral cells (MCs) from each glomerulus then transmit processed sensory patterns to higher order brain centers—such as mushroom bodies and lateral horn in insect brains or piriform cortex in mammals (Abel et al., 2001, Kirschner et al., 2006, Sinakevitch et al., 2017).

It has been well established in both insects (Fernandez et al., 2009, Locatelli et al., 2013, Chen et al., 2015) and mammals (Doucette et al., 2007, Doucette and Restrepo, 2008, Jones et al., 2008, Litaudon et al., 1997, Mandairon and Linster, 2009, Sullivan and Leon, 1986) that non-associative and associative plasticity in olfactory circuits shapes the output of activity patterns to odors. Presumably, this plasticity improves the contrast in neural representations required to help animals detect and discriminate important odors from background odors (Fernandez et al., 2009, Locatelli et al., 2016, Locatelli et al., 2013, Rath et al., 2011). This experience-dependent plasticity also implies that the neural representation of an odor could manifest differently in animals with different experiences. The glomerular activity patterns elicited by odors are canonically conserved across individuals of the same species and are established at the level of ORNs early in adult life prior to adult foraging in bees (Wang et al., 2005). However, experienced adults show slight deviations in the patterns of PN activation in glomeruli activated by the same odors (Arenas et al., 2012, Carcaud et al., 2012, Galizia and Kimmerle, 2004, Sachse and Galizia, 2002, Fernandez et al., 2009). This difference could arise as the AL develops while foraging as an adult to improve its ability to detect extant resource-associated odors as individual honey bees experience different floral resources (Chen et al., 2015, Fernandez et al., 2009). Honey bees freely fly in all directions within a several-mile radius of the colony to collect pollen and nectar resources from a diversity of floral sources that produce different perfumes to signal these rewards (Wright and Schiestl, 2009, Menzel, 1985). Alternatively, these mature adult differences in the AL’s PN responses to odors could represent random variability across individuals due to genetics or other developmental programming. To date, according to our knowledge, no one has experimentally investigated whether differences in individual experiences systematically influence odor processing in the brain, and to what extent experience contributes to inter-individual differences in perceptual abilities.

Chronological age is correlated with caste and experience in honey bees, and these factors affect the morphological development of the AL and mushroom body neuropils in the honey bee brain (Fahrbach et al., 2003, Fahrbach et al., 1995, Fahrbach et al., 1998, Farris et al., 2001, Brown et al., 2004, Brown et al., 2002, Coss et al., 1980). The mushroom body contains approximately 340,000 intrinsic Kenyon cells, which integrate inputs from visual, mechanosensory and taste modalities in addition to the olfactory inputs from the AL PNs (Rybak and Menzel, 1993, Strausfeld, 2002). Age and foraging experience have been correlated with region-specific volume changes and more complex dendritic arbors in the mushroom body calyx collar region, which receives inputs from visual neuropil (Durst et al., 1994, Fahrbach et al., 2003, Fahrbach et al., 1998, Farris et al., 2001). Additionally, foraging experience correlates with changes in the volume and synaptic density of glomeruli within the AL (Brown et al., 2004, Brown et al., 2002).

LNs play a modulatory role in the antennal lobe and synapse onto both other LNs and the AL output PNs. These neurons may be involved in the plasticity observed in the AL (Sinakevitch et al., 2011, Sinakevitch et al., 2013, Fernandez et al., 2009, Locatelli et al., 2013, Sachse and Galizia, 2002). It is speculated that some of this plasticity is modulated via octopamine (OA), which is critical to olfactory associative conditioning in both fruit flies, *Drosophila melanogaster*, (Gerber et al., 2009, Burke et al., 2012, Schwaerzel et al., 2003) and honey bees (Farooqui et al., 2003, Hammer and Menzel, 1998, Rein et al., 2013). In the honey bee AL, the octopamine 1 receptor (AmOA1) is expressed in GABAergic, inhibitory LNs and the AmOA1 expression patterns also exhibit inter-individual variability between experienced forager ALs (Sinakevitch et al., 2011). Thus it is possible that AmOA1 may be critical for the mechanistic steps leading to AL response differences between individuals honey bee foragers with different olfactory experience.

Previous studies have described inter-indiviual deviations in the glomerular representation of odors in the AL as well as several neuroanatomical features (Brown et al., 2004, Brown et al., 2002, Galizia et al., 1999a, Sinakevitch et al., 2011). In addition, acute odor exposures in the laboratory have shown that olfactory experiences modify the way in which odors are encoded in the AL (Chen et al., 2015, Fernandez et al., 2009, Locatelli et al., 2013, Rein et al., 2013). However we do not yet know the extent to which the olfactory experiences an animal receives while performing natural behaviors affect the modulation of olfactory circuits and the way odors are encoded and perceived. Toward those ends, we experimentally controlled the environmental complexity experienced by age-matched and genetically similar groups of forager honey bees and then directly measured inter-individual differences at morphological, physiological, and behavioral levels: expression of markers, including AmOA1, in the AL associated with neural plasticity and development, responses to odors in projection neurons in the AL, and odor mixture learning and recall, respectively. We show that natural olfactory experience significantly affects the tuning of olfactory responses in the honey bee AL and is necessary for adaptive AL odor processesing and odor-guided behavior in honey bees.

## RESULTS

### Experience-dependent maturation of the antennal lobe circuits in adult bees

The honey bee olfactory system is comprised of the antennae, the antennal nerve (AN), the antennal lobe (AL), and higher-order olfactory centers such as the mushroom body, lateral horn, and lateral protocerebral lobe (**Figure 1A)**. The AL consists of ~170 glomeruli, which is consistent with the number of olfactory receptor types in this species, and an aglomerular region (**Figure 1B**) (Robertson and Wanner, 2006, Sinakevitch et al., 2017). The glomeruli are divided into a cortex (or the outer region of the glomeruli) and a core region (Galizia and Sachse, 2010, Hildebrand and Shepherd, 1997, Nishino et al., 2009, Sinakevitch et al., 2017). The cortex region of the glomeruli contains arborizations of ORNs, PNs, and LNs and is located on the peripheral surface of each glomerulus (Nishino et al., 2009, Flanagan and Mercer, 1989b, Zwaka et al., 2016). The core region contains arborizations comprised of LNs and PNs and is located in the center of the glomerulus (Nishino et al., 2009, Zwaka et al., 2016, Sinakevitch et al., 2017).

**Figure 1.**
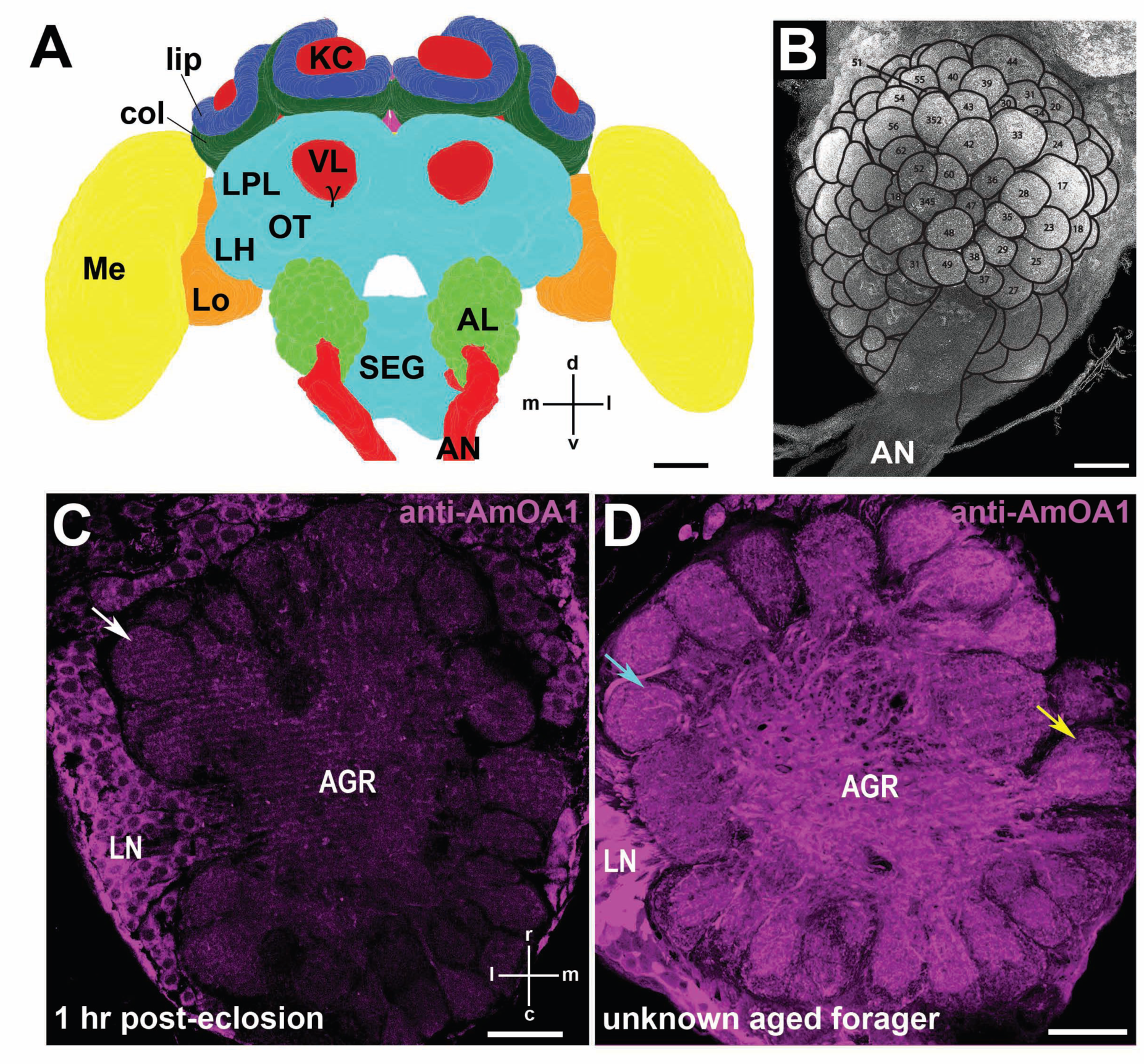
Honey bee brain and antennal lobe developmental plasticity with anti-AmOA1 stainings. **(A)** The schematic of the honey bee brain with major neuropilar areas. **(B)** Honey bee AL immunostained with anti-synapsin with an overlaid digital reconstruction of the AL identifying dorsal glomeruli that receive ORNS from (T1-T3 tracts). Glomeruli are identified and labeled according to Galizia et al. (1999a). Natural variation in anti-AmOA1 staining in a newly emerged adult bee (**C**) and an unknown aged forager (**D**). Arrows denote example glomeruli. Glomerulus outline in **D** highlights the core and cortex separations of yellow-arrow labeled glomerulus. Acronyms: KC: mushroom body Kenyon cells, lip: mushroom body lip region, col: mushroom body collar region; VL and γ: vertical lobe and gamma lobes of the mushroom body; LPL: lateral protocerebral lobe, OT: optic tubercle, LH: lateral horn, Me: medulla, Lo: lobula, AL: antennal lobe, SEG: Subesophageal ganglion, AN: Antennal nerve; d: dorsal, l: lateral, v: ventral, m: medial. Scale bar: **A**=250 μm, **B**= 50 μm, **C**= 20 μm, **D**= 20 μm

Newly eclosed adult workers show very little expression of the honey bee octopamine 1 receptor (AmOA1) within the glomerular (**Figure 1C**,—white arrow) or the aglomerular regions (**Figure 1C**). As a honey bee ages and begins performing foraging behaviors, there is a dramatic change in anti-AmOA1 receptor staining, with high level of intensity and heterogeneous expression within and across glomeruli in the glomerular and aglomerular neuropils of the AL (**Figure 1D**). For example, the two glomeruli marked by the teal and yellow arrows (**Figure 1D**) highlight this variation across glomeruli in one AL. In this forager the rostro-lateral glomerulus (teal arrow) is labeled by anti-AmOA1 in both the core and cortex. However in the medio-caudal glomeruli (yellow arrow), only the core is labeled by anti-AmOA1 (**Figure 1D**).

In order to explore the AL network changes that occur as a result of olfactory experience we placed bees in one of two olfactory experience treatments (**Figure 2**). We then immunolabeled the brains of foragers in our two treatment groups as well as newly emerged adult bees from the same cohort used to establish the treatment colonies. Since the AmOA1 receptor is known to be important for associative plasticity in the AL (Farooqui et al., 2004), we used anti-AmOA1 antibodies to study its distribution in the AL in relation to our treatments and age. Simultaneuously, we used Phalloidin-TRITC to visualize polymerized actin (F-actin) (Farris and Sinakevitch, 2003). High levels of F-actin staining are correlated with neurite growth and development and to the outgrowth of dendritic spines (Kaech et al. 1997; Groh et al. 2012).

**Figure 2.**
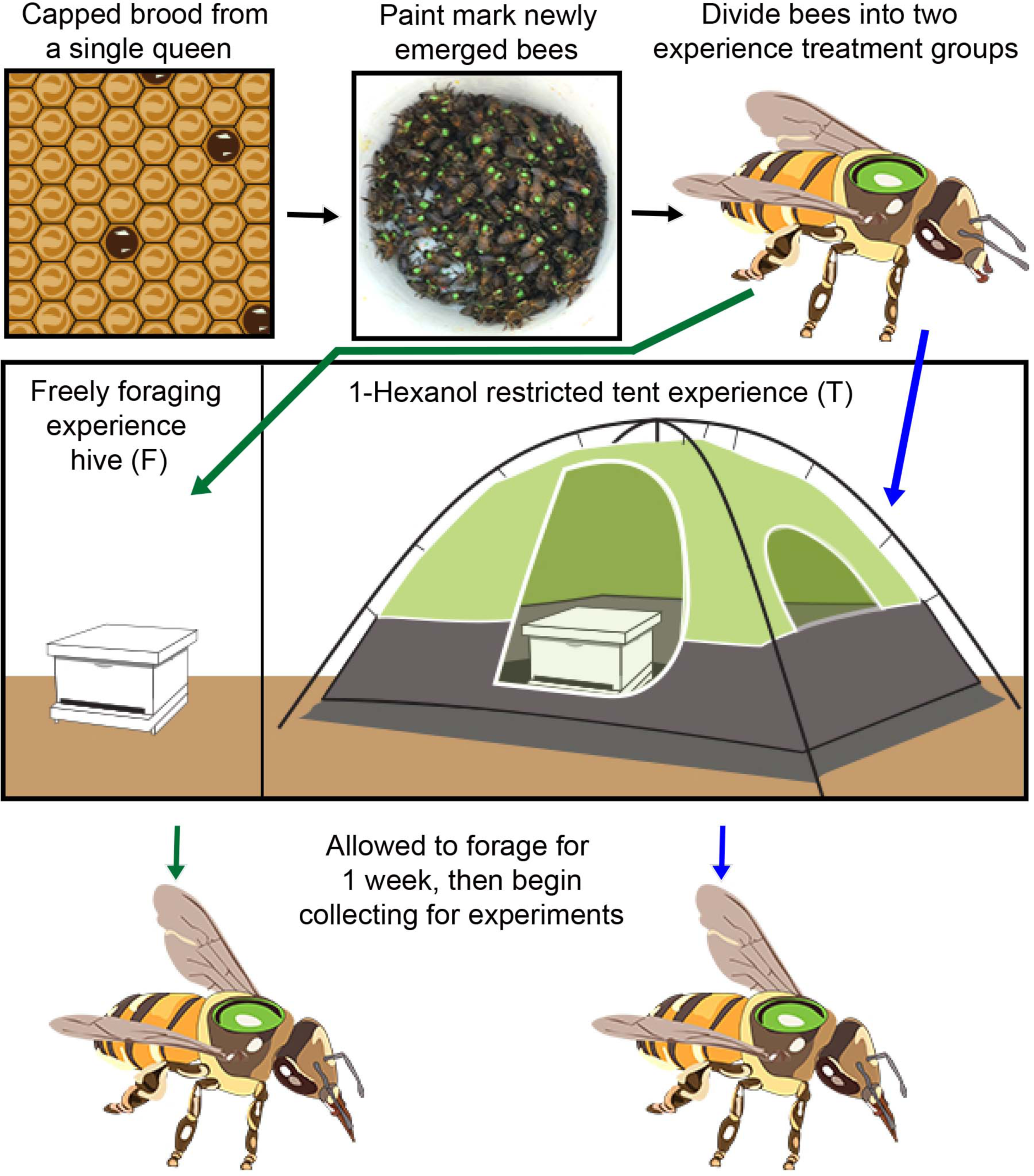
Experimental treatments for each experiment. Newly emerged honey bee workers collected from brood frame of a single colony and then paint-marked. Paint-marked workers were split into two treatment hives. Foragers from one hive were allowed to freely fly in the environment (F). Foragers from the second hive were restricted to forage inside of a tent (T) for pollen and sucrose resources, associated with a single pure odor 1-hexanol. After foragers had been observed foraging for 1 week, painted foragers were collected for calcium imaging at 31-40 days post eclosion, behavioral experiments (3-11 weeks post eclosion), and immunolabeling (34-40 days post eclosion).

To identify glomeruli in sectioned brains labeled with anti-AmOA1 and phalloidin, we first established a protocol using 3D reconstructions of the AL from whole mount preparations in which we identified glomeruli and made virtual sagittal sections (**Figure 3A)**. To ensure we were comparing similar synaptic regions across individuals, we only used saggital sections cut in a plane in which we could visualize both the T1 tract of the antennal nerve and distinct glomeruli, such as glomerulus 44 and its neighbors (**Figure 3B**). Using this method to identify glomeruli, we moved forward and stained sectioned ALs against both anti-AmOA1 and F-actin (**Figure 4**). We found a large shift in the expression patterns of anti-AmOA1 staining across AL development from newly emerged adult (**Figure 4A1,B1**) to aged forager (**Figure 4C1,D1**). In newly emerged adult bees, we primarily observed anti-AmOA1 staining in the cell bodies (white arrows) around the AL (**Figure 4A1,B1**). Staining was at low intensity across the aglomerular and glomerular regions. In contrast to anti-AmOA1 staining, phalloidin broadly stained F-actin in olfactory receptor neuron tracts and glomeruli at this stage of development (**Figure 4A2,B2**).

**Figure 3.**
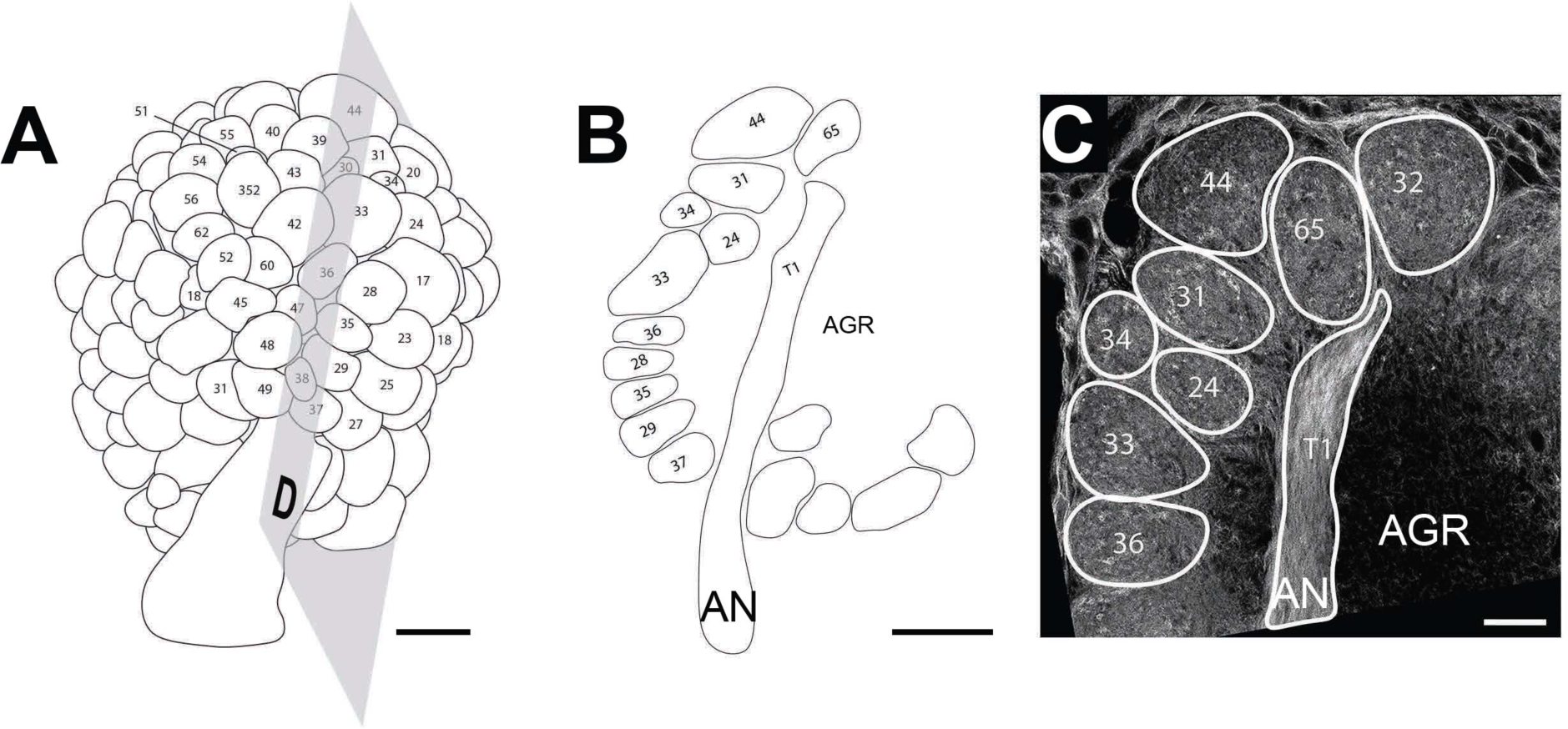
Antennal lobe glomeruli identification. **(A)** A digital reconstruction of the AL in (Figure 1B) Glomeruli are identified and labeled according to Galizia et al. (1999a). Shaded plane illustrates sagittal section displayed in B. **(B)** A digital, sagittal section taken from the reconstruction in A. Section clearly identified by the T1 ORN tract from the antennal nerve. **(C)** A digital overlay with labeled glomeruli and T1 ORN tract on sagittal sectioned AL. Tissue has F-actin labeled with Phalloidin-TRITC. AN: Antennal nerve, AGR: aglomerular region of the AL. Scale bar: **A**= 50 μm, **B**= 50 μm, **C**= 25 μm.

**Figure 4.**
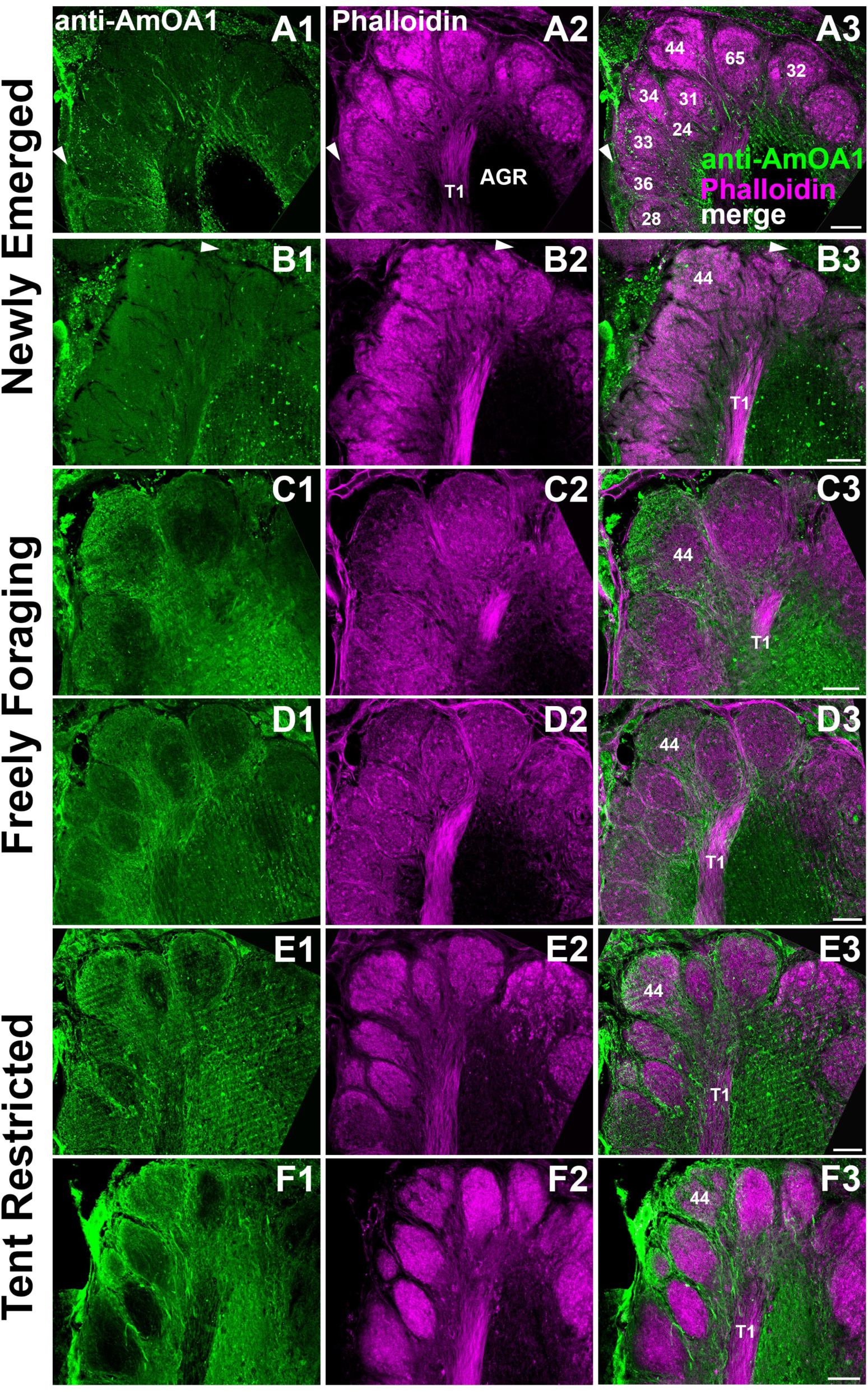
Sagittal sections of left AL from honey bee with different degrees of olfactory experience. Leftmost panel column shows anti-AmOA1 stainings (1), middle column shows F-actin bound by Phalloidin-TRITC (2), right-most panel shows merged images anti-AmOA1 and Phalloidin-TRITC labeling. **(A-B)** Newly emerged adult honey bee workers. **(C-D)** Adult forager honey bee workers aged 4-5 weeks post adult emergence. **(E-F)** Adult forager honey bees age-matched and sisters to C & D enclosed in a tent. Glomeruli are numbered in A3 and glomerulus 44 and T1 antennal nerve tract are labeled in all overlays for reference. Scale bars: 25 μm.

In aged, freely foraging adults, Anti-AmOA1 staining showed a more intense and variable staining pattern (**Figure 4C1,D1**). For example in figure C1, glomerulus 44 showed the high level of anti-AmOA1 staining in the cortex of the glomerulus, with much less staining within the core. We observed a different pattern for glomerulus 44 in a different bee in D1, which showed more intense anti-AmOA1 staining throughout both the core and cortex of glomerulus 44. Overall we observed a characteristically heterogeneous anti-AmOA1 staining across both the the aglomerular and glomerular regions as well as between glomeruli within tissue and between individuals (**Figure 4C1,D1**; Sinakevitch et al. 2011).

Freely foraging bees had a porous and sparse F-actin staining within glomeruli (**Figure 4C2, D2**). We primarily observed staining along the edges of the core and cortex of glomeruli and along olfactory neuron tracts, and to a much lower degree than observed in newly emerged adult bees. This pattern was highly consistent across preparations as exemplified in C2 and D2. Overall the merged images show very few fibers with close co-localization of AmOA1 and F-actin. However, technical limitations do not allow identification of AL cell types that express F-actin in glomeruli of these preparations.

In the tent-restricted treatment (T), we observed very similar anti-AmOA1 staining pattern to that found in bees with free-flying experience (**Figure 4E1,F1**). However, we observed significant differences in the F-actin stain. We observed a very high level of F-actin staining in both the core and cortex of all glomeruli, very unlike their naturally foraging, age-matched sisters (**Figure 4C2, D2**). Tent-restricted bees instead had staining patterns that were more consistent with those observed in newly emerged adult bees (**Figure 4E2, F2**) probably indicating still an yet immature condition.

### Experience-dependent effects on antennal lobe olfactory processing

To assess olfactory experience-dependent effects on AL odor responses, we visualized PN responses within glomeruli of the rostro-dorsal portion of the honey bee AL using a fura-2 dextran dye injection for age-matched bees that had one of our two experiential treatments (**Figure 2**). Using this calcium imaging method we then calculated the mean calcium responses in the PNs of each glomerulus in each bee during the odor stimulation period (**Figure 5**, see methods). A set of 14 different stimulations including different odorants, concentrations and mixtures were used to define the tunning profile of each glomerulus and evaluate if this tunning is affected by olfactory experience during adulthood. In addition, every stimulation with a given odor was repeated two times in each bee, thus we could also measure how consistent the response profile for a given glomerulus was within and across bees. The glomerular responses were highly consistent between the two presentations within bee in both treatments (Response profile correlations: F= 0.879 +/-0.028; T= 0.896 +/-0.0119). We next averaged these two presentations to get a mean glomerular response for each bee for each of the 23 identified glomeruli during the odor stimulation period (**Figure 5**). We then compared this mean odor response across bees. **Figure 6** shows the variability observed in the tunning of glomeruli 36 and 37 across free-flying bees at the level of output from the AL. The precise glomerular PN response profiles to the panel of odors were variable across bees (**Figure 6 A,B**). For example, glomerulus 36 responded consistently to 2-octanone (**Figure 6A**) and glomerulus 37 responded to acetophenone (**Figure 6B**) in all bees. However, the other odorants to which glomerulus 36 and 37 responded were variable between bees. Glomerulus 36 in bee 1 responded to both 1-hexanol and acetophenone in addition to 2-octanone. In bee 2, glomerulus 36 also responded to linalool and lemon oil (**Figure 6A**). Additionally, glomerulus 37 in bee 2 also responded to 1-hexanol, 2-octanone and linalool (**Figure 6B**). These inter-individual response deviations at the level of output from the AL (PNs) demonstrate differences in odor processing within the AL network between bees.

**Figure 5.**
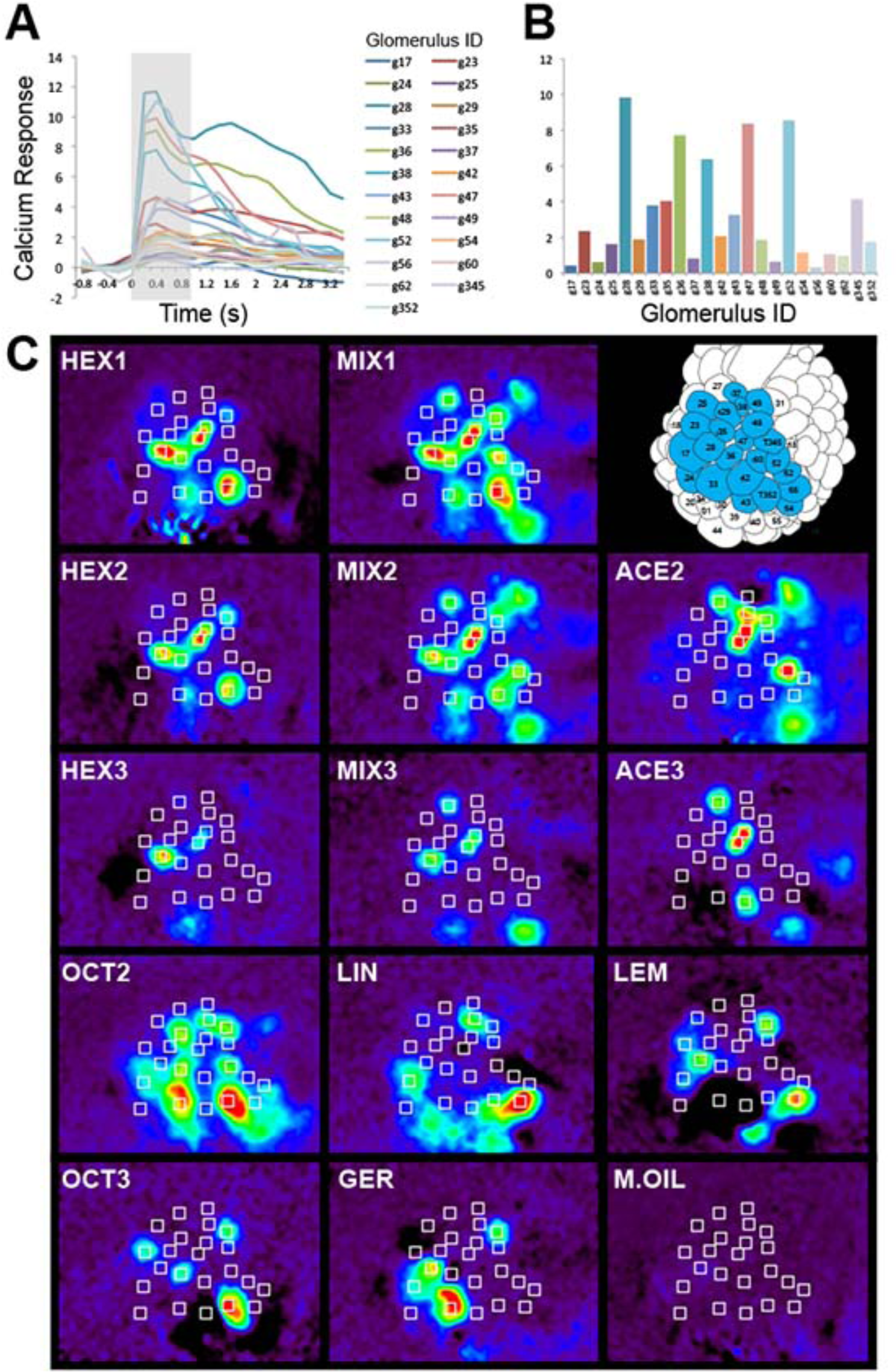
Calcium imaging data collection and processing protocol, example data from a single free-flying worker bee. (**A**) Response of 23 identified glomeruli to Mix1 over time. Shaded region denotes odor stimulation. (**B**) Averaged response during odor stimulation (shaded region in A) for each glomerulus. (**C**) Example false colored average response, over the 1 second odor stimulation window, of the honey bee AL to each odor stimulus presented. Schematic reconstruction of the rostral portion of the AL and the 23 identified glomeruli analyzed in this experiment in top right panel. Odors labels: HEX1 = 2×10^−2^ M 1-Hexanol, HEX2= 1×10^−2^ M 1-Hexanol, HEX3= 1×10^−3^ M 1-Hexanol, MIX1= 2×10^−2^ M 1-Hexanol and 1×10^−2^ M Acetophenone, MIX2= 1×10^−2^ M 1-Hexanol and 1×10^−2^ M Acetophenone, MIX3= 1×10^−3^ M 1-Hexanol and 1×10^−3^ M Acetophenone, ACE2= 1×10^−2^ M Acetophenone, ACE3= 1×10^−3^ M Acetophenone, OCT2= 1×10^−2^ M 2-Octanone, Oct3= 1×10^−3^ M 2-Octanone, GER= 1×10^−2^ M Geraniol, LEM= 1×10^−2^ M Lemon oil, LIN= 1×10^−2^ M Linalool. All odors dissolved in mineral oil (M.OIL).

**Figure 6.**
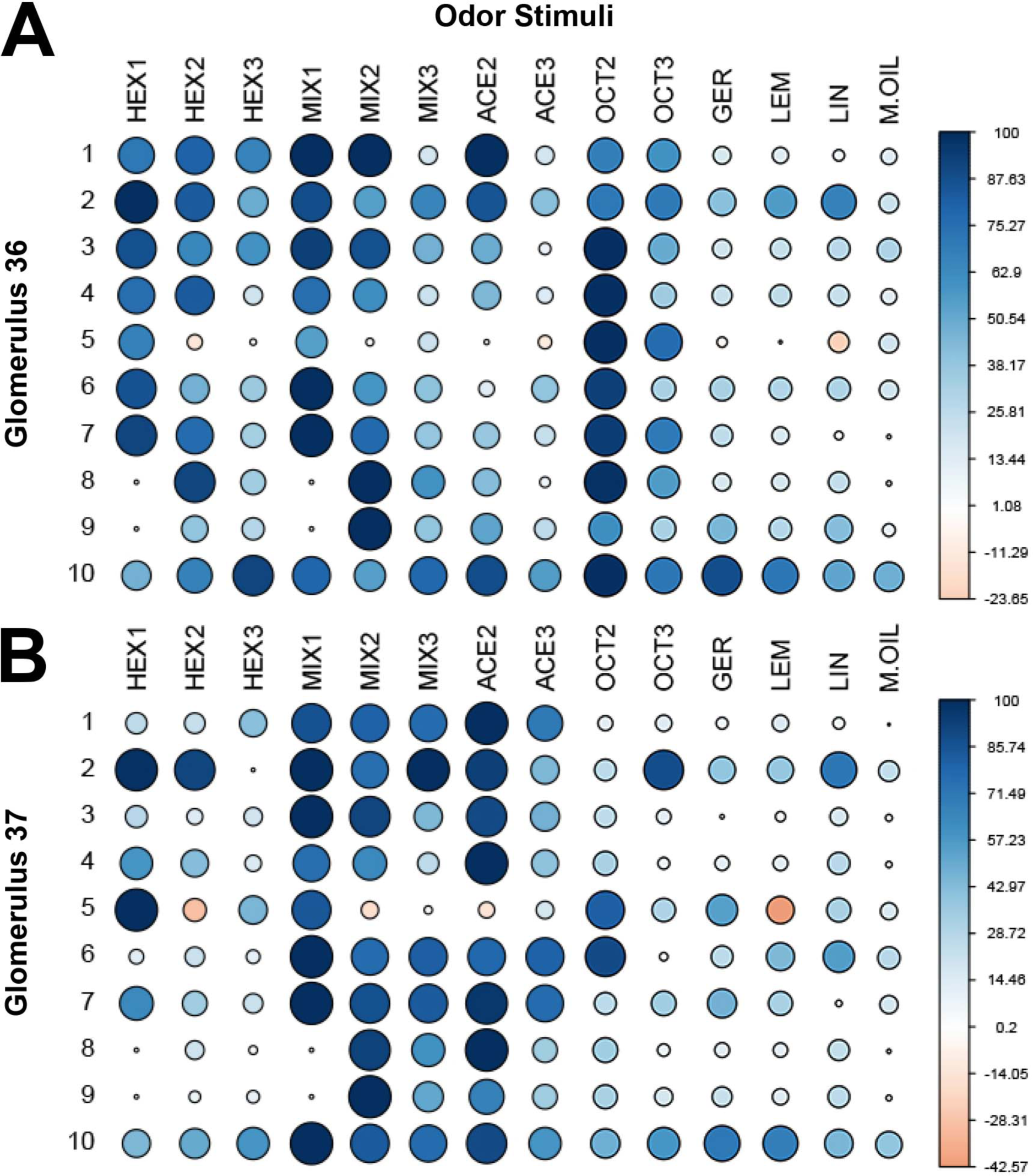
Natural inter-individual variation in glomerular response profiles to odors in free-flying honey bees. Response profiles for glomerulus 36 (**A**) and glomerulus 37 (**B**) for 10 age-matched foragers with natural olfactory experience. Size of circle denotes the magnitude of calcium response and and color the direction (excitation vs inhibition) of change. Odors labels: HEX1 = 2×10^−2^ M 1-Hexanol, HEX2= 1×10^−2^ M 1-Hexanol, HEX3= 1×10^−3^ M 1-Hexanol, MIX1= 2×10^−2^ M 1-Hexanol and 1×10^−2^ M Acetophenone, MIX2= 1×10^−2^ M 1-Hexanol and 1×10^−2^ M Acetophenone, MIX3= 1×10^−3^ M 1-Hexanol and 1×10^−3^ M Acetophenone, ACE2= 1×10^−2^ M Acetophenone, ACE3= 1×10^−3^ M Acetophenone, OCT2= 1×10^−2^ M 2-Octanone, Oct3= 1×10^−3^ M 2-Octanone, GER= 1×10^−2^ M Geraniol, LEM= 1×10^−2^ M Lemon oil, LIN= 1×10^−2^ M Linalool. All odors dissolved in mineral oil (M.OIL).

To test if this observed inter-individual variation in the glomerular response tunning was a consequence of dissimilar olfactory experiences across bees, we used a bootstrapping test of means to compare correlation matrices between bees within each treatment (see Methods). We first assessed odor-evoked response diversity of the individuals in each treatment group for all glomeruli. We found that there was a significantly higher correlation in the glomerular response profiles between the tent-restricted (T) bees than between the freely-flying (F) bees (All Glomeruli: paired t-test, p<0.001, Mean correlation value +/-SEM: Tent = 0.619 +/-0.027, Free = 0.545 +/-0.021). The increase of the correlation values indicates that the neural responses were less diverse across bees raised in the tent-restricted condition.

We next wanted to determine if the reduced inter-individual variation was driven by all of the 23 measured glomeruli or, alternatively, by only a few glomeruli that differ significantly between subjects. We compared the response profiles for a single glomerulus and repeated this comparison for each glomerulus between the two experiencial groups (**Figure 7**). Again, glomeruli for tent-restricted foragers tend to have lower response diversity (higher correlation) than the corresponding glomeruli in free-flying foragers (**Figure 7A**). This effect was significant in 12 of the measured 23 glomeruli (**Figure 7**, bootstrap test of means, 4 cases p<0.05, 8 cases p<0.01, R_Tent_<R_Free_ highlighted blue). We found 1 glomerulus with the opposite pattern, higher response diversity (lower correlation) in the tent-restricted compared to the free-flying foragers (**Figure 7,** bootstrap test of means, p<0.05, R_Tent_>R_Free_ highlighted red). Ten of the 23 glomeruli show no significant difference in the inter-individual variation between the two experiential treatments, versus 22 expected under the null hypothesis (**Figure 7** bootstrap test of means, p>0.05, R_Tent_ = R_Free_ highlighted grey).

**Figure 7.**
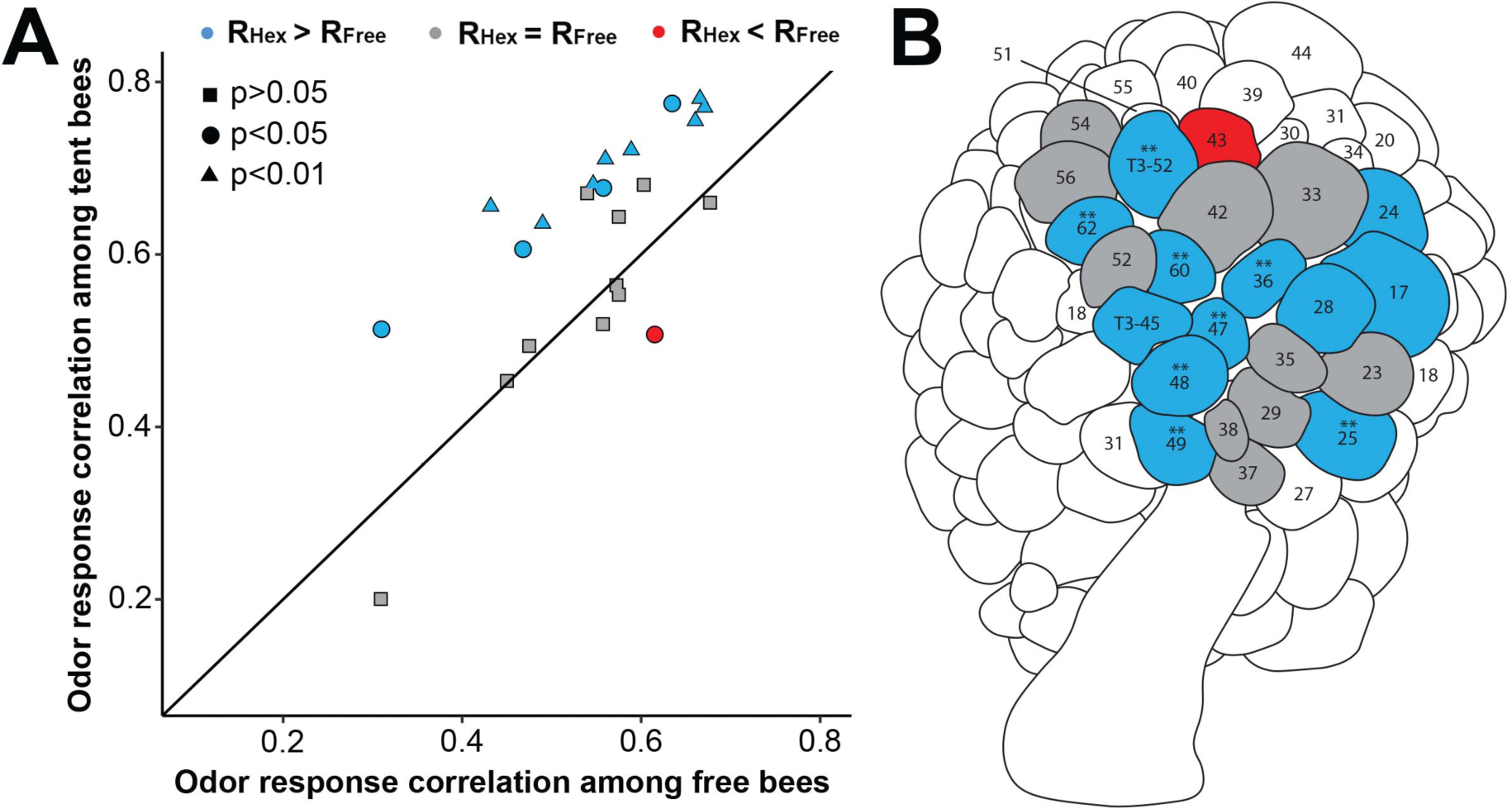
Increased correlation among the glomerular response profiles to odors in the tent-restricted treatment bees versus the free-flying bees. **A.** A comparison of the mean correlations among bees with a free-flying experience plotted against bees with a tent-restricted experience. Each point is the mean correlation for an identified glomerulus for the free-flying bees plotted against the distribution for the same glomerulus in the 1-hexanol tent-restricted experience bees. The diagonal line represents no difference in response space between conditions. This line represents the null hypothesis if the variation in odor-dependent glomerular responses between bees within each treatment are equal then all points should fall along this line. Colors denote significant differences between experience treatments and in which direction, the shape of the points denote the level of significance. **B.** Reconstruction of the honey bee AL and highlighted glomeruli denoting significant differences between experience treatments. Glomeruli with two stars above the number denote significance of p<0.01. Statistics: A bootstrapping test of means was used to assess significance and p-values were adjusted for multiple comparisons. Colors denote significant differences between experience treatments and in which direction, the shape of the points denote the level of significance.

### Experience-dependent effects on odor mixture learning

We next used a behavioral assay to compare the T (tent-restricted) and F (free-flying) odor experience groups to evaluate the effects of experience on odor learning. We chose a variant of an odor categorization problem reported in Wright and Smith (2004) in order to make the task more complex and thus more difficult to solve. This task required the animals to analyze a set of odor mixtures paired with reward, and then in a memory test determine if the animals were capable of generalizing from the mixture to the components. Animals experienced mixtures that were either ‘variable’ from trial-to-trial in their component concentrations or mixtures that remained ‘constant’ in their component across trials.

To control for odor specific effects, we also divided bees into two groups that received different mixture blends made up of 3 odorants (Mixture blend 1: acetophenone (target), geraniol, 2-octanone; or Mixture blend 2: phenylacetaldehyde (target), nonanal, α-farnesene). Each subgroup was further divided into bees receiving either a *variable* mixture training protocol (**Figure S1A**) or a *constant* mixture training protocol (**Figure S1B**). All groups received 16 presentations of the odor mixture paired with sucrose. Within the *variable* protocol, one odorant was held constant from trial-to-trial and was therefore the most reliable odorant signaling reward. This odorant is refered to as the ‘target’ odor. Bees were then tested for a proboscis extension response without reinforcement to each of the individual components that made up the associated mixture (**Figure S1A-C**; See methods and Wright and Smith, 2004 for further details).

Bees with a tent-restricted experience were able to associate mixture stimuli with sucrose just as well as free-flying adults over the 16 acquisition trials (**Figure S2**, GLM, family= Binomial, Experience*Blend: Z=1.389, p=0.165, Experience*Protocol: Z= −1.339, p=0.181, Experience*Blend*Protocol: Z= −1.762, p=0.078). As might be expected from the nature of the protocol, bees that experienced the variable odor mixture showed slower acquisition to the odor mixture than bees that received the constant protocol (**Figure S2**, GLM, family=Binomial, Variable protocol, Z= −4.702, p<0.001). We also found that bees with the tent-restricted experience associated odor mixture pairings more quickly than the free-flying bees irrespective of training protocol (**Figure S2**, GLM, family=Binomial, Experience, Z= 5.678, p<0.001). This difference could be due to a difference in sucrose response threshold, or perceived value, rather than a difference in learning ability (Ray and Ferneyhough, 1997). Tent-restricted bees were fed 50% sucrose ad libitum and the sucrose rewarded during training was also 50% sucrose, while the response threshold was unknown for free-flying bees that could have experienced higher quality nectar, both sugar content and other components, while foraging (Southwick et al., 1981, Alm et al., 1990).

We next tested for responses to all three components 2 hours later. According to prior work from Wright and Smith (2004), bees that were trained on the variable protocol should respond at a higher level to the component odors than those trained to the constant protocol, and they should also distinquish the target odor from the other mixture components. The target odor (acetophenone-Ace or phenylacetaldehyde-PAA) was held constant in each type of odor mixture during the variable training protocol. We found no difference between odor mixture blends (GLMM, family=Binomial, BeeID random factor, mixture blend*experience*protocol, t=2.877, df=458, p=0.3). As expected, the free-flying treatment bees that received the variable protocol responded more frequently to the target odor than bees trained using the constant protocol (**Figure 8A**), and the target odor elicited a higher response than the other components (**Figure 8A**; Figure S3A; GLMM, family=Binomial, BeeID random factor, Ace: t=2.544, df=457, p=0.01, PAA: t=2.256, df=457, p=0.025; non-target odors p>0.05). In contrast, the tent-restricted bees showed less difference in response to components beween the constant and variable protocols. (Figure 8B; Figure S3B, GLMM, family=Binomial, BeeID random factor, experience*protocol, t=-2.819, df=458, p<0.01). This difference in responses between the variable and constant training protocols across all odors was significantly larger in the free-flying bees than the tent-restricted bees across both mixture blends (Figure 8C, Mann-Whitney U test, W=1, p<0.01). Unlike their freely-flying counterparts, the bees with reduced olfactory experience did not distinguish the odor mixture components when trained to the variable protocol and instead responded at a high level across all odors irrespective of training protocol or odor blend.

**Figure 8.**
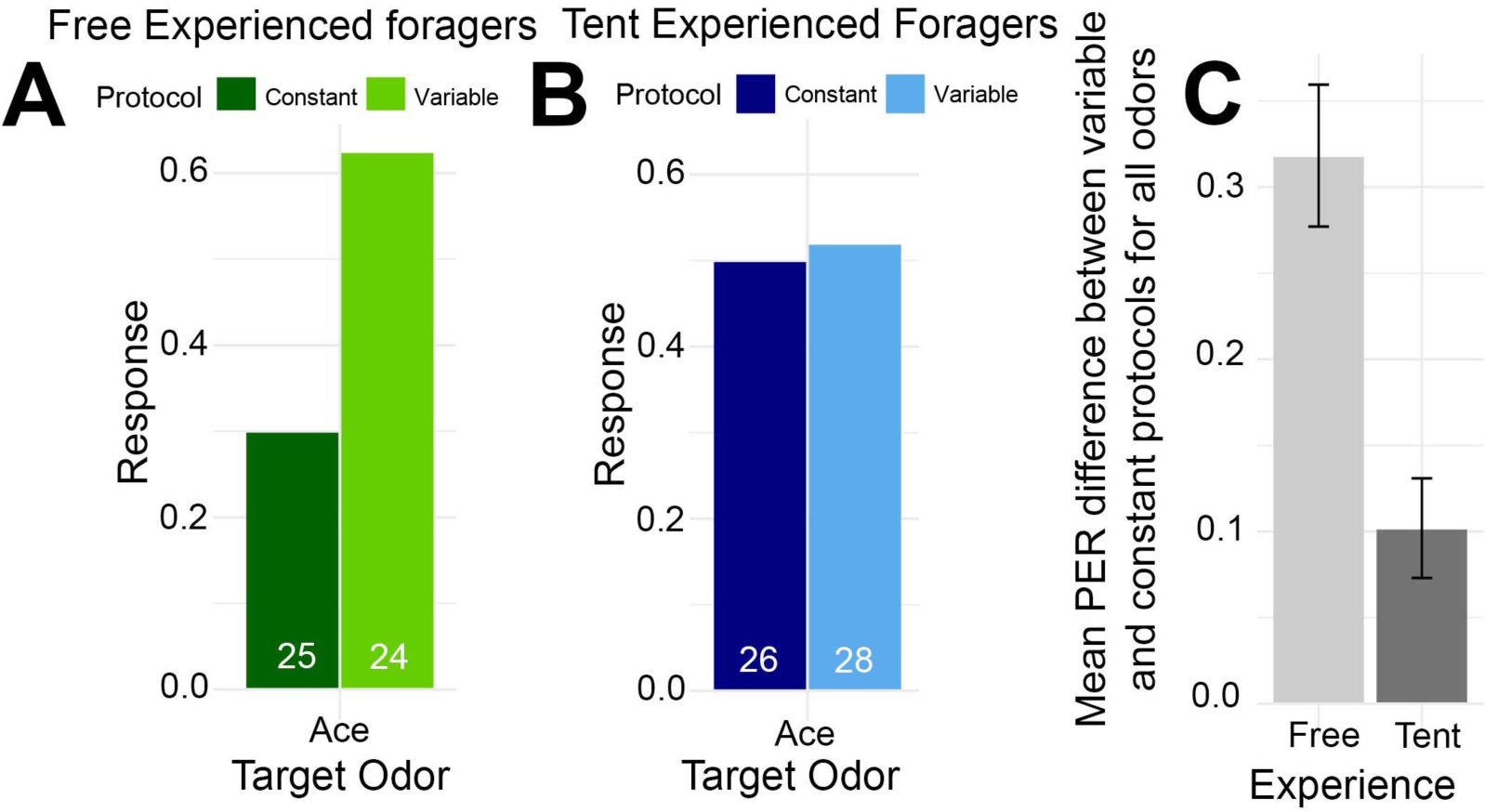
Bees were trained using either a constant mixture (dark color) or a variable mixture (light color) protocol. The bars show the proportion of bees extending their proboscis 2 hrs after conditioning to the target odor (acetophenone). See methods and Wright and Smith (2004) for more details. (**A**) Bees with a free-flying experience (green). (**B**) Bees with a tent-restricted foraging experience (blue). (**C**) Difference between the responses of bees trained using the variable mixture protocol and the constant mixture protocol for bees with a free-flying experience (light gray) and a tent-restricted foraging experience (dark gray) for all odor components in both trained mixture blends (See methods for details). Error bars represent standard error from the mean. Numbers at the bottom of each bar represent number of bees tested. Statistics: (**A-B**) GLMM with interacting factors of mixtures (blend1 or blend2), odor components (phenylacetaldehyde, nonanal, α-farnesene, acetophenone, geraniol, 2-octanone), bee experience (free vs. tent), training protocol (constant vs. variable), and bee identity as a random factor. There was no statistical difference in the responses to either mixture blend and just show responses to mixture blend1 target odor here, see Figure S2 for full odor response details. (**C**) Mann-Whitney U test, W=1, p <0.01, free constant N=45, free variable N=48, tent constant N=52, tent variable N=53.

## DISCUSSION

It has recently been shown that the mammalian olfactory bulb (OB) is constantly developing, even in mature adult animals (Gheusi et al., 2000, Nissant et al., 2009). Like the mammalian OB, there is a growing body of evidence that the insect antennal lobe (AL) is also continually tuned by experience. Like in mammals, short term exposure to an odorant has been shown to modulate the physiological responses to that odor (Mandairon and Linster, 2009, Locatelli et al., 2013). Responses to odors can also be modulated when presented in conjunction with reinforcing stimuli (Doucette et al., 2007, Mandairon and Linster, 2009, Sullivan and Leon, 1987, Fernandez et al., 2009, Chen et al., 2015). There is also evidence that the AL morphologically changes throughout adulthood in bees (Brown et al., 2004, Brown et al., 2002). This growing body of evidence suggests that, like the mammalian OB, a high level of plasticity in the insect AL continues through an animal’s lifetime. In this study, we show for the first time that naturalistic olfactory experience tunes the olfactory system during adulthood and quantify the extent to which olfactory experience affects the capabilities of an adult animal.

### Effects of olfactory restriction

Bees with natural foraging experiences may have foraged at multiple floral species, each of which has complex olfactory stimuli composed of many different volatile chemicals present at floral nectaries (Levin et al., 2003, Raguso, 2008). In our odor-restriction treatment, bees only had access to a single odorant consistently associated with the resources they collected. We have shown that by reducing the complexity of stimuli and controlling the olfactory experience of bees, there is an overall reduction in the inter-individual variation in how AL projection neurons respond to odors (**Figure 7C**). The chronic activation—or lack thereof—of specific glomeruli during sucrose reinforcement could have global effects on the synaptic weights of interneuron connections (LN-LN and LN-PN) in the AL, which could explain this result. This is supported by the high levels of F-actin observed in tent-restricted (T) bees, indicative of active structural or synaptic plasticity similar to that observed in newly eclosed bees (**Figure 4,** Phalloidin-TRITC) (Frambach et al., 2004, Groh et al., 2006). Furthermore, bees with a restricted experience responded similarly during recall to all the components of an associated mixture, even when receiving an association protocol that should produce strong responses to only one of the components (**Figure 8C**). This result is further supported by Cabirol et al. (2017), which recently found that experience deprivation results in a reduced ability to perform more complex learning tasks such as reversal learning.

### Pupal olfactory input on antennal lobe formation

There is a critical pre-eclosion period in AL development during which olfactory receptor neuron synaptic contact is necessary for both glomeruli formation and development in insects (Hildebrand et al., 1997, Oland et al., 1990). In brief, axonal projections from olfactory receptor neurons to the AL initiate the formation of preglomeruli (protoglomeruli in *Manduca*) (Hildebrand et al., 1979, Hildebrand et al., 1997, Oland et al., 1990, Groh, 2005). When animals are deprived of olfactory receptor neuron inputs during pupal development, via antennal amputation, the ALs develop without a defined glomerular structure (Hildebrand et al., 1979, Oland and Tolbert, 1987). With intact antennal connections and after preglomeruli formation, glial cells form a boundary that will later become the fully developed glomeruli (Oland et al., 1990, Oland and Tolbert, 1987, Oland and Tolbert, 1989). At this stage, but before each full synaptic glomerulus is formed, some PNs begin to arborize into their glomerular targets (Oland et al., 1990). The adult glomeruli can be visualized with anti-synapsin staining as the mature synapses form between the olfactory receptor neurons, local neurons, and projection neurons (Oland et al., 1990, Oland and Tolbert, 1989). More recently, it has been shown via genetic knockout of the obligatory co-receptor to all olfactory receptors of two ant species—*Harpegnathos saltator* and *Ooceraea biroi*—that normal development of the AL requires not only olfactory receptor neuron axon presence, but also responses to olfactory stimuli in those olfactory receptor neurons (Trible et al., 2017, Yan et al., 2017).

### Post-eclosion input on antennal lobe maturation

Post-eclosion experience shapes the development of neural networks in bees in the visual processing centers of the mushroom bodies (Coss et al., 1980, Fahrbach et al., 1995, Farris, 2005, Farris et al., 2001) and to some degree the early network development of the AL (Brown et al., 2004, Brown et al., 2002, Arenas et al., 2012). Wang et al. (2005) showed that global AL responses to odors, primarily composed of ORN responses, continued to develop up for up to two-weeks of age but prior to adult foraging began, which suggests that inputs to the AL develop and are stablizized prior to adult foraging. Fernandez et al. (2009), Locatelli et al. (2013), Hourcade et al. (2009), Arenas et al. (2009), Arenas et al. (2012), Chen et al. (2015) showed there are physiological changes in the AL of adult foragers in response to odors associated with different kinds of olfactory experiences, in which PN responses to odors are modified to support discrimination of relevant odors in AL ensemble activity.

Prior to the present work, in experiments with an experience-associated treatment, experimenters only isolated behavioral honey bee castes that did or did not have in-hive experience (Hourcade et al., 2010, Hourcade et al., 2009, Brown et al., 2004, Brown et al., 2002, Coss et al., 1980, Fahrbach et al., 1995, Farris, 2005, Farris et al., 2001), gave general odor experience in the hive for a fixed period of time (Arenas et al., 2009, Arenas et al., 2012) or measured age-groups independent of caste or experience (Wang et al., 2005). They then measured physiological changes and/or region-specific morphological changes in volume and synaptic densities (Coss et al., 1980, Fahrbach et al., 1995, Farris, 2005, Farris et al., 2001, Brown et al., 2004, Brown et al., 2002, Arenas et al., 2009, Arenas et al., 2012, Wang et al., 2005). In the present study, we extend prior studies by both allowing animals to develop within the hive normally and test animals from the same genetic cohorts and within the same caste that have different olfactory experiences. We also for the first time add a more comprehensive view of the effects of olfactory restriction at the level of neuroanatomy, physiological responses to odors, and a complex odor recall assay.

Our data provide a more comprehensive picture of the post-eclosion, experience-dependent effects of odor processing in bees. In light of previous work, our data suggest that experience post-eclosion is necessary for the production and/or maintenance of a mature AL network that is capable of performing complex olfactory-based tasks. We do not yet know the identity of the specific neuronal sub-types and network connections affected during olfactory restriction. However, restriction likely affects more than one sub-population of neuron within the AL and could possibly also affect inputs from higher-order brain regions.

In our treatments, we manipulated the chronic food-odor associations that adult honey bee foragers had access to. The tent-restricted bees had access to all of the normal colony odors. Thus, the primary olfactory difference between groups was specifically the diversity of food-associated odors that they experienced. However, we introduced additional differences when we restricted bees’ activities to a tent. In addition to olfactory experience, we manipulated the general foraging experience of bees by restricting flight distance and aerobic effort, visual diversity, nectar complexity, and likely the colony forager dance communication profile. Most of these effects likely have an influence on other brain regions as well as the antennal lobes.

### Perception of complex odor mixtures

The differences in recall of odor mixtures by bees with restricted vs. unrestricted experience could have several possible explanations that are worthy of further study. First, bees with limited experience could have difficulty recognizing odors in general. These bees could also have less precise recall of odor mixtures compared to bees with free-flying experience, or they could have a higher degree of generalization to odor associations. These possibilities could be tested using a discrimination task across a number of odorants followed by a recall test to similar and dissimilar odorants as done in Guerrieri et al. (2005). Coding of complex olfactory mixtures is not well studied in the AL, despite these stimuli being abundant in natural settings (Laloi et al., 2000, Strutz et al., 2014, Locatelli et al., 2016, Guerrieri et al., 2005). More complex olfactory stimuli would likely be computationally more intensive to process; therefore, it is likely that if the AL were somehow underdeveloped (as our data suggest), processing mixtures could be more difficult for odor-restricted bees compared to their natural foraging counterparts.

### Summary

Our data show that adult olfactory experience affects the AL network in honey bees. We show that the development of the AL neuropil is delayed, the inter-individual variation in the physiological responses to odors in this network is reduced, and the capacity of animals to learn and behaviorally analyze odor-mixtures is significantly reduced when animals are deprived of natural *food-associated* odor experiences as adults. These findings strongly suggest that experience drives the normal inter-individual variation we observe in nature and that these olfactory experience are necessary for the normal functions of the AL in bees.

## METHODS

### Beekeeping and rearing

A single open-mated honey bee queen (*Apis mellifera carnica*) was caged on an empty frame for 2 days and allowed to lay eggs on a single frame. All projeny from an open-mated queen will be full-or half-sisters depending on whether they share a paternal genotype or not (Page, 2013). After egg laying, the queen was released to move freely in the hive. Just prior to adult emergence, the frame was placed in an incubator. Newly emerged (eclosed) adult honey bees were collected from the frame within 24 hours of eclosion and were paint marked with a non-toxic pen (Sharpie, oil based). A total of 1000 newly emerged bees were marked and split into 2 groups of approximately 500 bees every 2-4 weeks. Each of these groups of ~500-700 of bees was introduced into one of two 10-frame queenright host hives, named by T and F. We used one pair of host hives for both the calcium imaging and immunohistochemical experiments, and two pairs of host hives for the behavioral experiments.

The hives were placed within 30 meters of one another in a shaded courtyard. Hive T was enclosed by an individual 15.9 m^3^ (8×10×7ft) mosquito net tent in which all of the sucrose and pollen resources were artificially provided. The second hive (F) was left outside of the tents and foraged freely at the Tempe, AZ campus of Arizona State University. (**Figure 2**).

The food resources in the tents were marked with a single-odor stimulus 1-hexanol (hive T) as an artificial CS. This was done as follows: The 50% sucrose (w/w) solutions provided to the tent-restricted bees contained 0.01% (w/w) odor. Around both the sucrose and pollen resources, a 10% odor solution diluted in mineral oil was applied to an absorbent material placed around both resources. Sucrose solutions were replaced every 24 hours and a new 10% odor mixture was placed around the resources once in the morning and once in the afternoon. To the human nose, all materials retained the smell of the odor, even after 24 hours.

### Honey bee collection

Paint-marked honey bee foragers were all collected when returning to the hive from a foraging event approximately 7 days after the first paint-marked bee was observed foraging.

#### Immunostaining collection

Paint-marked bees from all hives were collected simultaneously over three days and immediately processed for for fixation with the following immuostaining procedures. These bees were between 34-40 days old from adult emergence.

#### Calcium imaging collection

Paint-marked bees were collected from all hives simultaneously over a three-week time frame and were immediately harnessed and prepared for calcium imaging. The honey bee cohort used in experiments were switched from the original 1000 newly emerged bees to a cohort 2-4 weeks younger half way through the experiment, in order to keep the length of foraging experience more consistent across bee subjects. All bees were between 31-40 days old from adult emergence.

#### Variance learning PER assay collection

Approximately 7500 paint-marked bees were collected between the months of Dec. 2016 and June 2017, paint-marked newly emerged bees were placed into each hive at regular 3-4 week intervals. Age groups were switched approximately every 3-4 weeks, depending upon the dominant foraging paint marked cohort at the time of collection. Eight bees were collected from each hive experience treatment (T, tent-restricted, and F, free-flying) each training day and they were divided equally into each training protocol group (see PER assay below).

### Calcium Imaging

#### Bee preparation and in vivo PN staining

Marked bees were captured, briefly cooled on ice and restrained in custom made individual holders suited for calcium imaging (Galizia and Vetter, 2004). After recovery from cooling, the bees were fed with 1.0 M sucrose solution and left undisturbed until staining shortly after. A window was cut in the top of the head capsule, dorsal to the joints of the antennae and rostral to the medial ocellus. The hypopharyngeal glands and trachea near the alpha-lobes (Rybak and Menzel, 1993) were moved and served as visual reference for the staining (Sachse and Galizia, 2002). The tip of a glass electrode coated with fura2-dextran (potassium salt, 10.000 MW, ThermoFisher Scientific) was inserted into both sides of the protocerebrum, dorsolateral to the vertical-lobes, aiming for the lateral antennalprotocerebral tract (l-APT) that contains the axons of uniglomerular PNs (Galizia and Rössler, 2010). A few seconds later, after the dye had dissolved we closed the window in the head capsule using the piece of cuticle that had been previously removed. The dye was left to travel along the l-APT tracts until the next day, roughly 10-18 hours later. Before imaging, the antennae were fixed pointing toward the front, where odor will be delivered, using a low-temperature melting wax Eicosane. Body movements were prevented by gently compressing the abdomen and thorax with a piece of foam held in place by a piece of tape. The brain was then rinsed with Ringer’s solution (130 mM NaCl, 6 mM KCl, 4 mM MgCl2, 5mM CaCl2, 160mM sucrose, 25mM glucose, 10mM HEPES, pH 6.7, 500 mOsmol; all chemicals from Sigma-Aldrich), and glands and tracheae covering the ALs were removed. When necessary we also cut a small hole between the antennae and mandibles, and then pulled out a small section of a compact structure of muscles, esophagus, and supporting chitin. We did this to put this structure under slight tension and pull it away from the brain to prevent accessory movements in the AL (Mauelshagen, 1993). Only ALs that presented homogenous staining of all visible glomeruli were used for imaging. Stained bees were then mounted on the microscope and were allowed to recover for 15 minutes before imaging. We collected 18 full recordings, 10 recordings from bees that received the free-flying treatment and 8 bees that received the tent-restricted treatment

#### Data acquisition and analysis

A Polychrome V (Till-Photonics, Gräfelfing, Germany) was used to emit excitation at two wavelengths alternating between 340 and 380 nm. Imaging data was then collected at 5 Hz using a CCD camera (SensiCamQE, Till-Photonics) mounted on an upright fluorescence microscope (Olympus BX-50WI, Japan) with a 20x objective NA 0.95 (Olympus) using a 505 DRLPXR dichroic mirror and a 515 nm LP emission filter (Till-Photonics). The final spatial resolution of each image was 1376 × 1040 pixels with a pixel side length equaling 2.6 μm. The exposure times during excitation were 8 ms at 340 and 2 ms at 380 nm. The image analysis was done using custom software written in the Interactive Data Language (IDL; Research Systems) using routines created by Giovanni Galizia (University of Konstanz, Germany). Measurements from each animal consisted of a sequence of 50 fluorescence images, obtained at each excitation wavelength (F_*i*_^340^, F_*i*_^380^, where subscript *i* is the number of images 1 to 50, and superscript denotes measurements at the excitation wavelengths 340 nm or 380 nm). Calcium responses were calculated as the ratio R_*i*_ = (F_*i*_^340^/F_*i*_^380^) × 100. We subtracted the background responses (R_*b*_) from these ratios. We calculated R_*b*_ by averaging the R_*i*_ values 1 second immediately before the odor onset, where R_*b*_ = 1/5 (R_11_+ …+ R_15_). The resulting relative calcium response measure (ΔR) represents a percentage change from the odor-free reference window (R_11_–R_15_). This measure has previously been shown to be directly proportional to changes in intracellular calcium concentration (Galizia and Kimmerle, 2004). Next, we identified glomeruli based upon their morphology and relative position using our own AL reconstructions and the digital atlases of the honey bee AL (Flanagan and Mercer, 1989a, Galizia et al., 1999a). We also visualized glomeruli using the raw fluorescence images obtained at the 380 nm excitation wavelength. For an additional confirmation of glomeruli locations we created images that represent the degree of correlation between neighboring pixels with a tool provided by Mathias Ditzen (Freie Universitaet Berlin, Germany). Pixels stemming from the same glomerulus are highly correlated over time and pixels from different glomeruli are not. We finally ended up with a common set of 23 glomeruli that could be identified across all animals. All glomeruli corresponded to the dorso-rostral side of the AL innervated by the antennal nerve T1 tract: glomeruli 17, 23, 24, 25, 28, 29, 33, 35, 36, 37, 38, 42, 43, 47, 48, 49, 52, 54, 56, 60, 62; and the T3 tract, 54, 52 (Flanagan and Mercer, 1989a, Galizia et al., 1999a). The activity for each glomerulus was calculated by averaging mean ΔR activity over a 9×9 px square area that corresponds to approximately a 23.4 × 23.4 μm square which fits within the center of each glomerulus. We then averaged this activity for each glomerulus in each animal over the 1-second odor stimulation period. These values were then used for the final comparisons of odor-elicited activity across animals and experience treatments.

In order to quantitatively measure the degree of inter-individual variation within each treatment we created correlation matrices to compare bees within each treatment group (Tent N=8, Free N=10)., We considered two scenarios: A) The odor-response variation observed across individuals in a given glomerulus, and B) The same variation but pooled across all glomeruli. For A), each of the cells in the correlation matrices were the Pearson correlation value of the measured mean odor-evoked glomerular responses to all tested odors, as described above, between two individual bees within the same treatment group considering each glomerulus in a different correlation matrix (23, 8×8 Tent matrices and 23, 10×10 Free matrices). We used a bootstrapping test of means to compare the correlation matrices between the two treatment groups. All p-values were adjusted for multiple comparisions according to the False Discovery Rate (Benjamini and Hochberg, 1995). For B) each of the cells in the matrix was the correlation pooled across all glomeruli between two individual bees within the same treatment group (one, 8×8 tent matrix and one, 10×10 Free matrix). We used a paired t-test to compare the responses of pooled glomeruli.

#### Odor stimulation and imaging session

The focus of this analysis was to determine whether a given glomerulus was recruited or not by a given odor, and if this glomerular response profile varied across animals within a given treatment group. For that aim, the glomerular responses were measured for pure odors, mixtures and different concentrations of both. Odors were diluted in mineral oil: 1-hexanol 2×10^−2^, 1×10^−2^ and 1×10^−3^; acetophenone 1×10^−2^ and 1×10^−3^; mixture1 (1-hexanol 2×10^−2^ + acetophenone 1×10^−2^); mixture2 (1-hexanol 1×10^−2^ + acetophenone 1×10^−2^); mixture3 (1-hexanol 1×10^−3^ + acetophenone 1×10^−3^); 2-octanone 1×10^−2^ and 1×10^−3^; geraniol 1×10^−2^; lemon oil 1×10^−2^ and linalool 1×10^−2^ M. Ten μl of odor solution were loaded onto a filter paper strip (0.5 × 4 cm) that was put into a 1 ml glass syringe, which served as an odor cartridge. The odor-delivery device had 14 identical channels, each composed of a three-way solenoid valve (LFAA1200118H; The LEE Company) and an odor cartridge. Valve opening was synchronized with the optical recordings using the acquisition software TILLVisION (Till-Photonics). When the valve opened, the air volume inside the cartridge was delivered (~50 ml/min) into a continuous charcoal filtered air stream (~500 ml/min) which in turn directed the air toward the honey bee head. Thus, the final concentration of odors reaching the honey bee was actually ~1/10 of the concentration in the headspace of the cartridge. Imaging acquisition trials lasted 10 seconds and were separated from each other by one minute. Odor stimulation lasted one second and started three seconds after onset of acquisition. Each odor was tested two times in each animal, making a total of 28 stimulations, including blank trials with mineral oil. Odor order was randomized, with the only restriction being to not use the same odor in two consecutive trials. Behind the honey bee, an exhaust continuously removed air, keeping the arena clean of olfactory stimuli.

### Immunocytochemistry

After collection, bees were immobilized by cooling on ice for a maximum of three minutes, the heads were cut from the abdomen and placed into the 4% paraformaldehyde in phosphate buffer saline (PBS) pH 7.4, and the brains were removed and placed in 1 ml of fixative overnight at 4 °C.

#### Primary antibodies

Affinity-purified goat polyclonal anti-AmOA1 antibodies (21^st^ Century Biochemicals, Inc. Marlborough, MA) were raised against a synthetic peptide acetyl-AMRNDRSPSYSMQVPQQGC-amide, which corresponds to amino acids 547-564 of the honey bee AmOA1 receptor. These antibodies were previously used to study the distribution of the AmOA1 receptor in the honey bee brain (Sinakevitch et al., 2011, Sinakevitch et al., 2013). Mouse monoclonal anti-synapsin antibodies (SYNORF1, 3C11) were purchased from DBH (Data Bank Hybridoma). Anti-synapsin binds to protein associated with presynaptic sites of neurons and is largely used for labeling the synaptic neuropil. Phalloidin conjugated with TRITC (Tetramethylrhodamine Isothiocyanate (Invitrogen) binds to polymerized actin (F-Actin) and was used to label areas of high synaptic turnover and localize areas where growth of (active) synapses occurred.

#### Secondary antibodies

We visualized primary anti-AmOA1 using F(ab’)2 fragments of donkey anti-goat antibodies conjugated to Cy5 (Jackson ImmunoResearch Laboratories). F(ab’)2 fragments of donkey anti-mouse antibodies conjugated to Alexa 488 were used to visualize anti-synapsin (Jackson ImmunoResearch Laboratories).

### Anti-AmOA1 staining procedures on brain sections

Fixed brains were washed in phosphate buffered solution and embedded in 8% (w/v) agarose (low melting point, Sigma) in water. The 80 μm sections of brains were made using a vibrating blade microtome (Leica VT1000S, Leica Biosystems, Germany) in PBS. Sections were washed (6×20 minutes) in 0.5% of Triton X-100 in PBS (PBSTX) and then were pre-incubated with normal donkey serum (Jackson ImmunoResearch Laboratories) for 15 minutes. Next, the primary antibodies goat anti-AmOA1 in the PBSTX was added to brain sections at a 1:16 dilution for overnight incubation at room temperature. Next, sections were incubated overnight at room temperature with secondary antibodies (donkey anti-goat antibodies conjugated with Cy5 at 1:200) and Phalloidin-TRITC. Sections were then washed in PBS (6×10 minutes) and embedded on slides in mounting medium. The specificity and control tests for the goat anti-AmOA1 stains are described in detail in prior work (Sinakevitch et al., 2011, Sinakevitch et al., 2013). In the current study, we peformed additional control tests to demonstrate that neither the primary goat anti-AmOA1 antibodies nor the secondary anti-goat Cy5 antibodies interacted with the Phalloidin staining. In the first control, the primary AmOA1 antibodies were omitted from the protocol and we added the secondary antibodies as described above. In the absence of the primary AmOA1 antibody we observed no differences in the Phalloidin staining patterns compared to Phalloidin stained alone preps. Additionally in the reverse control, omitting the anti-goat Cy5 secondary antibodies there was no difference compared with the Phalloidin staining observed [data not shown]. This indicated that neither the primary goat anti-AmOA1 antibody nor the secondary anti-goat Cy5 interacted with the conjugated Phalloidin staining.

### Anti-synapsin staining procedures on whole-mount brains

After fixation procedures described above, brains of free-flying honey bees (*Apis mellifera carnica*) were washed in PBSTX (6×20minutes). Following washes, brains were pre-incubated with normal donkey serum for 15 minutes and incubated for three nights at room temperature with anti-synapsin at 1:800 in PBSTX solution. The following day, brains were washed again in PBSTX (6×20minutes) and incubated for three nights at room temperature with donkey anti-mouse antibodies conjugated with Alexa 488 at 1:270 and then with Phalloidin-TRITC at 1:160 in PBSTX overnight. After staining with secondary antibodies, brains were washed with 4% paraformaldehyde fixative for 10 minutes and then put through a dehydration protocol using increasing steps of ethyl alcohol. Following full dehydration and three washes in 100% ethanol, brains were cleared in methyl salicylate. They were then mounted on slides in methyl salicylate.

### Confocal Image collection and processing

Images were collected using a Leica TCS SP5 confocal laser scanning microscope (Leica, Bensheim, Germany) with a Leica HCX PLAPO CS 40x oil-immersion objective (numerical aperture: 1.25) using the appropriate filter and laser setting for each florescent molecule used (see sections above). Image stacks were collected using 1μm optical sections and a total volume of approximately 20 μm. Image stacks were then flattened using maximum intensity functions in the Leica software. Image size, intensity, and resolution were adjusted using Adobe Photoshop CC.

### 3D reconstructions of the antennal lobe and whole brain

To identify the glomeruli number in sectioned preparations we used AVIZO software (Thermo Scientific, FEI) to make a 3D reconstruction of the AL from a whole mount preparation. We then identified glomeruli by comparison with the honey bee brain atlas (Rybak, 2012), Flanagan and Mercer (1989a) and Galizia et al. (1999a), and made virtual (sagittal) sections of this reconstruction. We also reconstructed the whole brain of one honey bee forager for general reference. Tiff files of whole mount brain confocal scans were taken every 10 μm throughout the right and left honey bee AL recontsructions and scans were taken every 2 μm for the whole brain reconstruction. These files were imported into AVIZO software, and the voxel dimension outputs of each image stack were imported to provide correct dimensions (Thermo Scientific, FEI). Using the image segmentation function, we identified brain structures based upon the honey bee brain atlas and identified individual glomeruli on the dorsal surface of the left and right ALs (Flanagan and Mercer, 1989a, Galizia et al., 1999a, Rybak, 2012). Brain neuropils and glomeruli were individually labeled by hand throughout the image stacks. A volume rendering of these labels was created, and exported into Photoshop CC. In Photoshop, we created a manipulatable 3D image using the 3D volume function.

### Proboscis extension response variance learning

Eight harnessed bees from each odor experience treatment group (F and T) were collected and presented with an odor mixture paired with 50% sucrose (w/w) at a 10-minute inter-trial interval using an odor delivery system each training day (Smith and Burden, 2014, Wright and Smith, 2004). Stimuli were delivered by passing air through a glass cartridge containing 20 μl of an odor mixture on a small strip of filter paper. Each odor cartridge was used for no more than six presentations.

Odor blends of three monomolecular odorants were used and mixed with hexane as a solvent. Two classifications of odor mixtures were produced. The first blend was comprised of acetophenone, geraniol, and 2-octanone. The second blend consisted of phenylacetaldehyde, nonanal, and α-farnesene.

As in Wright and Smith (2004), in the variable mixtures, one of the three odors was presented consistently over all trials at a constant concentration. Other odors were varied from trial to trial, using either a high (H: 2.0M) or low (L: 0.0002M) concentration (Wright and Smith, 2004). Four mixtures were prepared for each odor set as follows: LLH, LHL, LLL, and LHH. With the target odor, phenylacetaldehyde or acetophenone, presented consistently at the low concentration for each blend. Ratios of each odor mixture were then confirmed using GC/MS. It is important to note that the total mean odor concentrations across all presentations were equal for both protocols. Each subject was presented the above odor mixtures over 16 trials in a pseudorandomized order, with each odor being presented four times over the process of training.

An additional group of bees was trained in the same way as above, however, the odor mixture ratios were held constant across the entire training process. Bees from both the natural free-flying experienced group and the tent-restricted odor experience group were trained under either the varied or constant mixture conditions. At the end of conditioning, the bees were placed in a humidified container. After 2 hours, the subjects were tested for a PER response to the individual component odors that comprised the training odor mixtures, separately, at the low concentration (0.0002M). Responses to each odor component were presented without any sucrose reinforcement, and PERs were recorded (Wright and Smith, 2004). Test odors were presented in a randomized order.

We analyzed responses for acquisition using Generalized linear model (GLM), and for recall using Generalized Linear Mixed Model (GLMM) to allow bee identity to be included as a random factor. Finally, we compared differences in recall responses between the variable and constant acquisition protocols between treatment groups using the non-parametric Mann-Whitney U test.

## ACKNOWLEDGEMENTS

This work was supported by awards from NIGMS (GM113967) and NSF IOS (1556337) to BHS. RCG was supported by NIMH (1R01MH106674) and NIBIB (1R01EB021711). FFL was supported by CONICET (National Scientific and Technical Research council), Argentina. We would also like to thank Drs. Ronald Rutowski, Jon Harrison, Stephen Pratt, Jason Newbern, and Karla Moeller for their comments on this manuscript.

