## Supplemental Figure 1 for "Experience-dependent tuning of early olfactory processing in the adult brain"

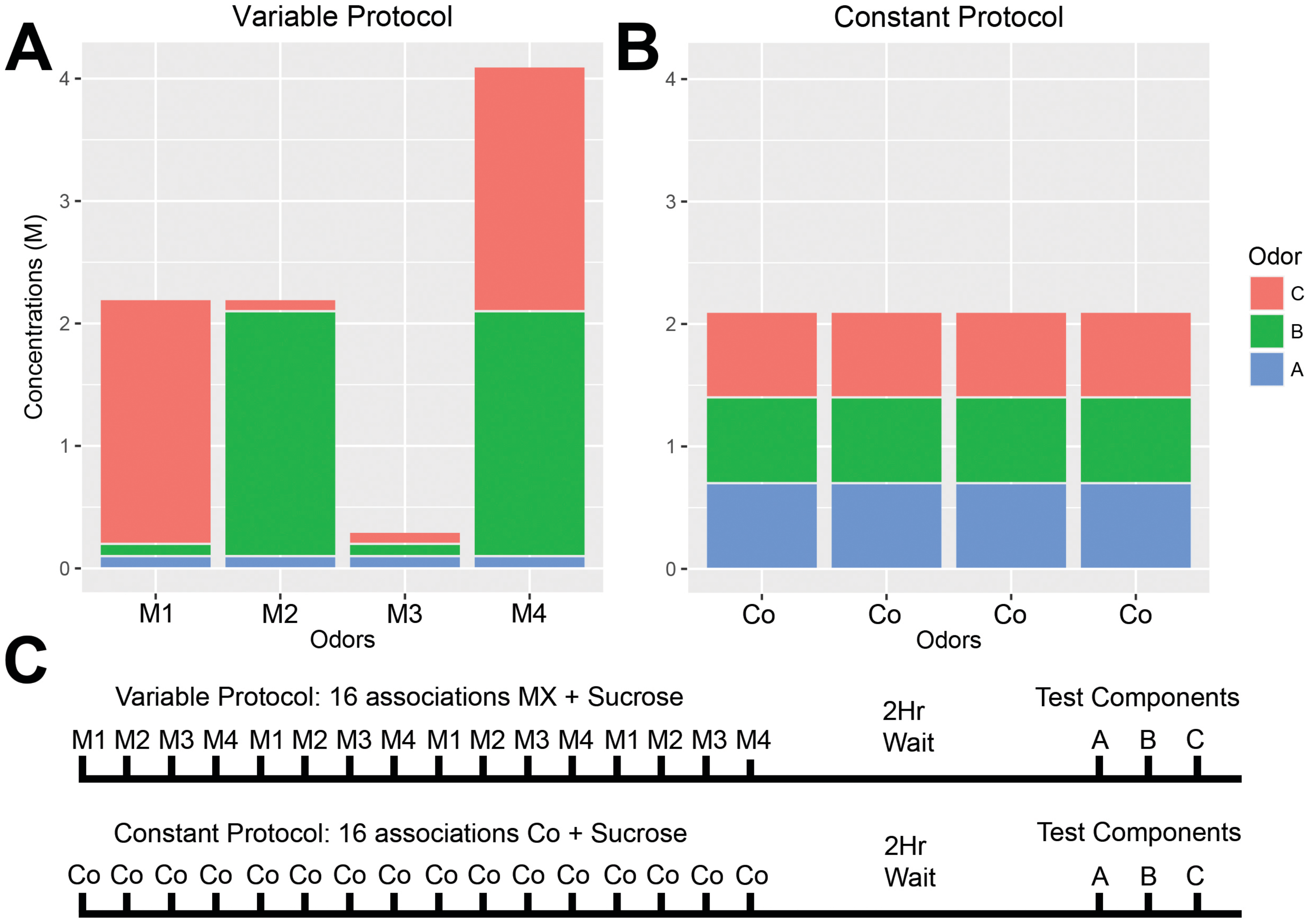


**Figure S1.** Performance of honey bee foragers aged 4-11 weeks post adult emergence. The odor mixture component concentrations used during the (A) variable mixture protocol (B) the constant mixture protocol. (C) A schematic of the associative conditioning and component memory testing procedures. Bees that received the variable protocol received different mixtures (M1, M2, M3 or M4) in a pseudorandomized order over 16 associative acquisition trials. Bees that the constant protocol received constant mixtures (Co) over 16 associative acquisition trials. Mean total odorant exposure was equivalent across the two protocols. Memory test component order was randomized between bees for both protocols. We used a mixture blend made of three components: A=Acetophenone, B=Geraniol, and C=2-Octanone. Acetophenone (A, the target odor) is held at a constant intensity even in the variable protocol. Across trials, the average mixture intensity is the same for both the constant and the variable protocols. We also tested a second mixture blend see methods and Wright and Smith (2004) for more details.
