## Supplemental Figure 2 for "Experience-dependent tuning of early olfactory processing in the adult brain"

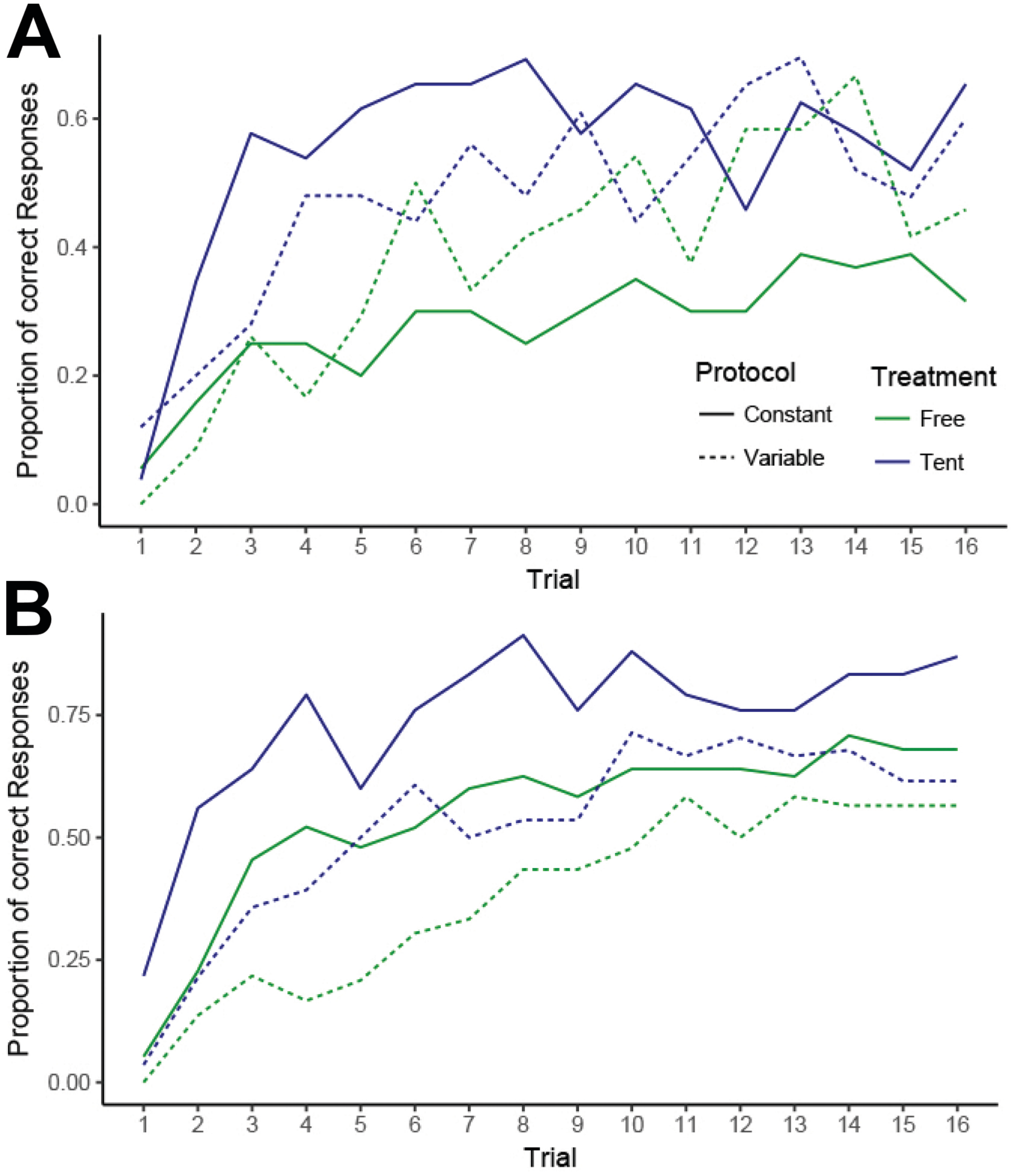


**Figure S2.** Acquisition of odor mixture blend 1 across freely foraging (green) and tent bees (blue) which received either a constant (solid line) or a variable (dashed line) mixture protocol. (A) Mixture blend 2 odors (B) mixture blend 1 odors. Sample Sizes: free blend 2 constant N=20, free blend 2 variable N=24, free blend 1 constant N=25, free blend 1 variable N=24, tent blend 2 constant N=26, tent blend 2 variable N=25, tent blend 1 constant N=26, tent blend 1 variable N=28.
