## Supplemental Figure 3 for "Experience-dependent tuning of early olfactory processing in the adult brain"

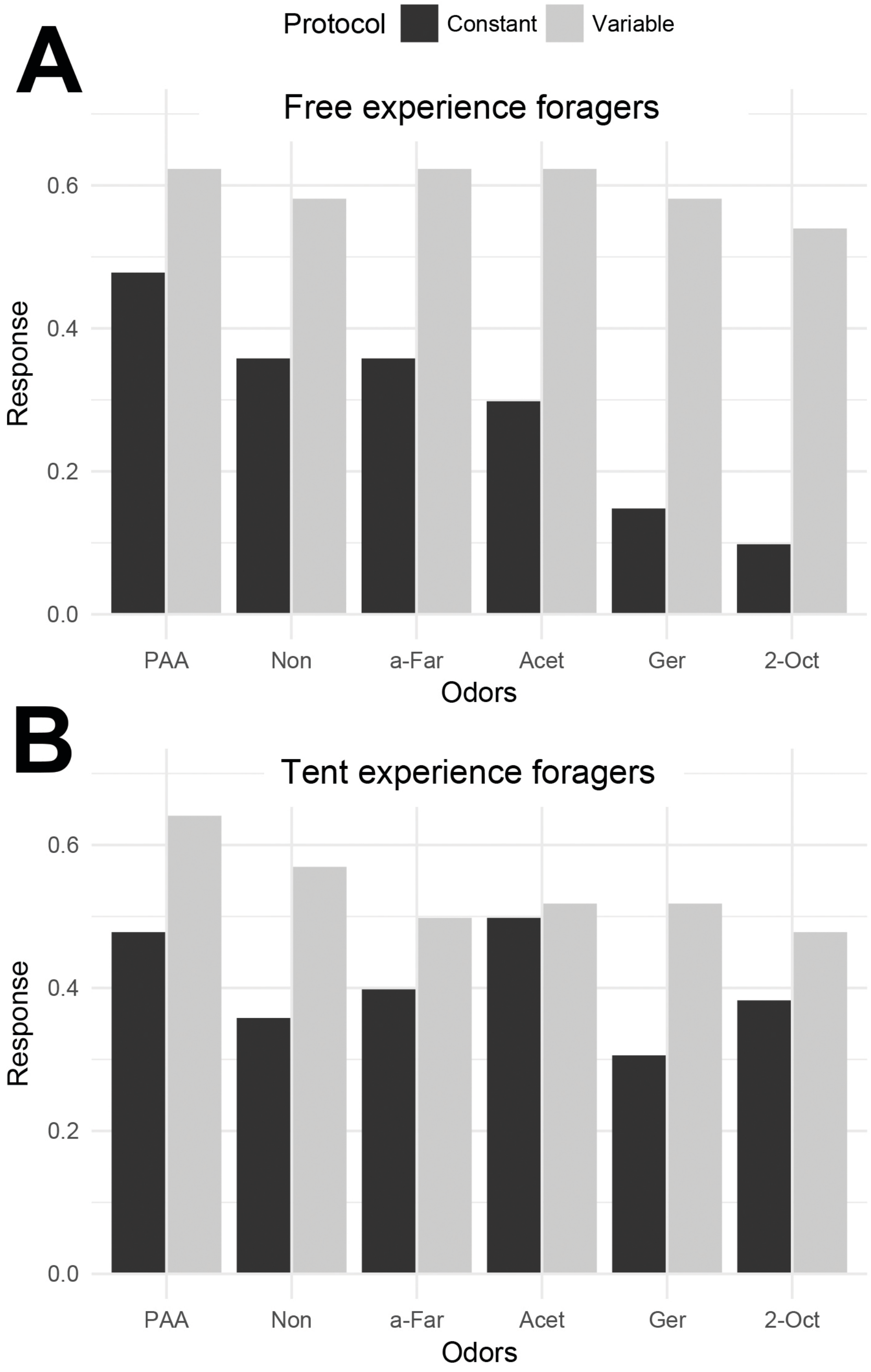


**Figure S3.** The memory response rates for bees 2hrs after receiving the constant acquisition protocol (black) or the variable acquisition protocol (gray) for bees with (A) a free-flying experience and (B) a tent-restricted foraging experience. Sample Sizes: free blend 2 constant N=20, free blend 2 variable N=24, free blend 1 constant N=25, free blend 1 variable N=24, tent blend 2 constant N=26, tent blend 2 variable N=25, tent blend 1 constant N=26, tent blend 1 variable N=28.
